# Assessing the effect of model specification and prior sensitivity on Bayesian tests of temporal signal

**DOI:** 10.1101/2024.08.12.607579

**Authors:** John H Tay, Arthur Kocher, Sebastian Duchene

## Abstract

Our understanding of the evolution of many microbes has been revolutionised by the molecular clock, a statistical tool to infer evolutionary rates and timescales from analyses of biomolecular sequences. In all molecular clock models, evolutionary rates and times are jointly unidentifiable and ‘calibration’ information must therefore be used.

For many organisms, sequences sampled at different time points can be employed for such calibration. Before attempting to do so, it is recommended to verify that the data carry sufficient information for molecular dating, a practice referred to as evaluation of temporal signal. Recently, a fully Bayesian approach, BETS (Bayesian Evaluation of Temporal Signal), was proposed to overcome known limitations of other commonly used techniques such as root-to-tip regression or date randomisation tests. BETS requires the specification of a full Bayesian phylogenetic model, posing several considerations for untangling the impact of model choice on the detection of temporal signal. Here, we aimed to (i) explore the effect of molecular clock model and tree prior specification on the results of BETS and (ii) provide guidelines for improving our confidence in molecular clock estimates.

Using microbial molecular sequence data sets and simulation experiments, we assess the impact of the tree prior and its hyperparameters on the accuracy of temporal signal detection. In particular, highly informative priors that are inconsistent with the data can result in the incorrect detection of temporal signal. In consequence, we recommend: (i) using prior predictive simulations to determine whether the prior generates a reasonable expectation of parameters of interest, such as the evolutionary rate and age of the root node, (ii) conducting prior sensitivity analyses to assess the robustness of the posterior to the choice of prior, and (iii) selecting a molecular clock model that reasonably describes the evolutionary process.

**Author summary:** Our knowledge of when historical and modern pathogens emerged and spread is largely grounded on molecular clock models. The inferences from these models assume that sequence sampling times must have captured a sufficient amount of evolutionary change, which is typically determined using tests of temporal signal, such as BETS. Although BETS is generally effective, here we show that it can incorrectly detect temporal signal if the chosen evolutionary model makes implausible statements about the evolutionary timescale, a situation that is difficult to diagnose, particularly with complex Bayesian models. We demonstrate that this problem is due to a statistical artefact, that we refer to as tree extension and that it can be minimised by conducting careful prior predictive simulations, and by eliciting biologically plausible priors in the model. Overall, our study provides guidelines for improving our statistical confidence in estimates of evolutionary timescales, with key applications for recently emerging pathogens and data sets involving ancient molecular data.

## Introduction

Molecular sequence data have been essential to unravel the evolutionary history of many organisms. Furthermore, phylogenetic methods are particularly useful and have a wide range of applications, from resolving the timing of evolutionary relationships [1], to quantifying differential abundance testing and mediation analyses [2, 3].

The molecular clock is a statistical approach to estimate the date of phylogenetic divergence events based on the hypothesis that molecular evolution, in the form of substitutions, follows an identifiable statistical process. Under the earliest and simplest molecular clock model, known as the strict clock, substitutions are assumed to accumulate at a constant rate over time and across lineages [4] (for an in-depth introduction to molecular clocks see [1]). At the other end of the spectrum, relaxed molecular clock models allow every lineage in a phylogenetic tree to display a different evolutionary rate [5] (reviewed in [6]).

All molecular clock models have a fundamental limitation; that evolutionary rates and times are jointly unidentifiable. That is, there exist an infinite number of combinations of evolutionary rates and times that are compatible with a given amount of evolutionary divergence [7, 8]. For this reason, external information allowing us to constrain some of the parameters of the model must be used, a process known as a molecular clock calibration. The finding that some organisms accumulate substitutions in a measurable timescale prompted the use of sequence sampling times for calibration, a practice known as ‘tip calibration’ [9, 10]. The latter is particularly useful for microbial organisms for which fossil information cannot be used, or is not available, to constrain internal node dates.

A fundamental requirement for using tip calibration is that the sequenced data were sampled from a measurably evolving population, i.e. that the interval of time over which the samples were taken captures an appreciable amount of evolutionary change in the studied organism [11]. For rapidly evolving microbes, such as RNA viruses, this might already be achieved by drawing samples over weeks or months. For more slowly evolving microbes sampling over many years or centuries may be needed. Importantly, whole genome sequencing has been a boon for molecular clock analyses of slowly evolving microbes because the resulting data sometimes contain enough information to warrant calibration using sequences collected over a few months or years [12].

There exist several statistical tests to determine whether a sampled population has measurably evolving behaviour, also known as tests of temporal signal. The root-to-tip regression takes a phylogenetic tree for which the branch lengths measure evolutionary distance (i.e. a phylogram) and fits a linear regression of the distance from the root to the tips as a function of their sampling time [10]. The regression slope is a crude estimate of the evolutionary rate, the *x-*intercept is the time to the most recent common ancestor, and the *R*^2^ is a measure of clocklike evolution. In general, the root-to-tip regression is a powerful tool for visual inspection of the data, for example to detect outliers or identify lineages with particularly low or high evolutionary rates [13–16]. However, because the data points are not statistically independent, resulting statistics such as *p-*values are invalid and cannot be used as formal statistical tests of temporal signal [17]. A different approach, known as the ‘date randomisation test’, consist of fitting a molecular clock to the data after permuting the sampling times multiple times to obtain a ‘null’ distribution of the evolutionary rate [18]. The data are considered to have temporal signal if the evolutionary rate estimated with the correct sampling times falls outside such ‘null’ distribution [18–20].

Although date randomisation tests can be computationally efficient [21], they make it difficult to exploit the full extent of Bayesian phylodynamic models. For example, it is not clear how to incorporate uncertainty in sampling times. A more powerful alternative that is fully Bayesian is the Bayesian Evaluation of Temporal Signal (BETS) [22]. The premise of this test is that the data sampled from a measurably evolving population should have higher statistical fit when the sampling times are included than when they are not, which can be assessed through model selection. In practice, the data are analysed with their correct sampling times (i.e. heterochronous) and with all samples assigned the same date (i.e. isochronous, with the sampling time set to the same point in time), while keeping the rest of the phylogenetic model the same, including the molecular clock, tree prior and substitution model. For clarity, an isochronous tree is one where the sampling times are identical, also known as an ‘ultrametric’ tree (where the distance from the each of the tips to the root is identical) [22, 23], and thus a heterochronous tree is one where the sampling times are different.

The log marginal likelihood is calculated in each case to compute log Bayes factors, which quantify the amount of evidence for one model over another, here that with sampling times vs that without. When considering log Bayes factors, a value of 3 or greater is deemed ‘strong’ evidence, while a value below is deemed as ‘not worth more than a bare mention’ [24]. Further, any value greater than and including 5 is considered ‘very strong’ evidence. A major advantage of BETS is that it can consider the full model and it can accommodate important sources of uncertainty, such as that from radio carbon dating of ancient DNA samples [25] (reviewed in [26, 27]).

Most parameters of the phylogenetic model have individual prior probability distributions that can be chosen by the user, for example, the evolutionary rate, or the transition-to-transversion ratio of the HKY substitution model. The prior for the phylogenetic tree topology and branch lengths is usually a branching model, such as a coalescent or birth-death process, which implicitly imposes a prior probability distribution on the ages of nodes, and therefore may inadvertently conveys highly informative calibration priors. Moreover, model selection, as used for BETS, can be sensitive to the choice of prior, even if the posterior is not [28, 29]. Here we investigate the impact of the tree prior and is associated parameters, and the molecular clock model in the detection of temporal signal. We also explore alternative parameterisations of the full Bayesian model that can improve the accuracy of tests of temporal signal.

## Results

### Empirical data analyses

We first explored the effect of model specification on the results of BETS using empirical data sets for which temporal signal was previously detected using other methods. The following three data sets were used: *Vibrio cholerae* [30], the bacterium responsible for cholera; *Powassan virus (POWV)* [31], a tick-borne virus; and *Treponema pallidum* [32], the bacterium that causes syphilis. The *V. cholerae* and *T. pallidum* data sets include ancient samples, and the phylogenetic trees from the POWV and *T. pallidum* indicate complex population structure that is typical of data sets from multiple outbreaks.

We conducted BETS analyses under a coalescent tree prior with constant population size and two possible clock models; a strict and an uncorrelated relaxed clock with an underlying log-normal distribution [5]. Our choice of the constant-size coalescent tree prior is based on statistical convenience, as it is fully parametric, but it is not necessarily an accurate representation of the biological process. We set up our analyses in BEAST1.10 [33] and calculated log marginal likelihoods with and without sampling times for each combination of molecular clock model and tree prior (for a tutorial of using BETS see: https://beast.community/bets_tutorial).

To assess the impact of the tree prior we set different prior distributions for the effective population size, *θ*, the only parameter in the constant-size coalescent (sometimes the priors on parameters of the tree prior are known as a hyperpriors [34]). In the exponential-growth coalescent, which we also considered in our simulations (see below), the equivalent parameter is known as the ‘scaled population size’ (denoted with the Greek letter Φ) and it is proportional to the population size at present [35]. Both, *θ* and Φ are referred to as a *scale parameters* for time because large values imply more dispersion (the molecular clock rate is also a scale parameter), and they are typically assigned a 1*/x* prior distribution, which is the Jeffrey’s prior that is uninformative and invariant to reparameterisation [36]. The 1*/x* prior has attractive attributes because it maximises the signal from the data, but it is an improper distribution (it does not integrate to one over its domain, because 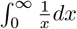 is undefined), a problem for model comparison using Bayes factors, because marginal likelihood calculations require that all priors be proper distributions [37, 38]. Instead, we selected three prior distributions, an exponential, Γ (Gamma), and log-normal, that have been used in recent literature [32] (this Γ prior on the scaled population size is the default in the BEAST1.10.5), and as shown in Table 1 (for the epidemiological or demographic interpretations of the parameters of the tree prior we refer the reader to [36, 39]).

**Table 1.** Prior distributions for the effective population size of the constant-size coalescent (known as *θ* in the constant-size coalescent and different to the scale parameter of the Γ distribution).

| Probability distribution function | Parameters |
| --- | --- |
| Exponential | mean, $\mu = 1.0$ |
| Log-normal | mean, $\mu = 1.0$; standard deviation $\sigma = 5.0$ |
| $\Gamma$ (Gamma) | shape, $\kappa = 0.001$; scale, $\theta = 1000$ |

Our rationale for using different prior distributions on *θ* is its impact on parameters that pertain to the molecular clock. In particular, under the coalescent process, the expected time of divergence between sampled lineages is inversely proportional to the population size, meaning that large values of *θ* will result in an older time to the most recent common ancestor than small values for this parameter. The prior on *θ* will also have an impact on the evolutionary rate for two key reasons. First, by impacting the overall age of the tree, it impacts the length of time over which the sequence data evolved. Second, the default prior for the evolutionary rate in BEAST1.10 is a Γ distribution with shape (*α*) of 0.5 and beta (*β*, also known as the ‘rate’) equal to the tree length (sum of all branch lengths) [40, 41]. In this software, this prior is known as the CTMC-rate reference prior and its mean value is 0.5*/*tree length [41–43], meaning that it is indirectly impacted by *θ* (or Φ).

To illustrate the interactions between parameters in the posterior expression for a full Bayesian phylogenetic model, consider expression 1, where *T* is the phylogenetic time-tree, *θ* is the effective population size of the coalescent, *κ* denotes the parameters of the substitution model (e.g. the transition-to-transversion ratio of the HKY), *r* is the evolutionary rate under a strict molecular clock model, and *D* is the sequence data (for an example using plate notation see [44]). In this model, the data are treated as heterochronous, such that *r* is estimated, the tree prior is a constant-size coalescent, the prior on the evolutionary rate is a CTMC-rate reference prior, and the substitution model is a HKY.

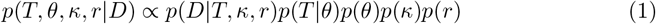

The posterior probability *p*(*T, θ, κ, r*|*D*) is proportional to the phylogenetic likelihood, given by *p*(*D*|*T, r, κ*), multiplied by the remaining terms, collectively known as the prior. The tree prior, *p*(*T*|*θ*) is sometimes known as the phylodynamic likelihood [45], and here it is the probability of the tree given *θ*. Note that *p*(*r*) ∼ Γ(0.5, *tree length*) (the CTMC-rate reference prior), implying that *p*(*r*) is also dependent on *T* and therefore *θ*, via *p*(*T* |*θ*).

The *V. cholerae* data set displayed overwhelming support for temporal signal (Table 2 and Fig 1), regardless of the molecular clock model and prior on *θ*, with log Bayes factors of over 200. Note that a log Bayes factor of 3 corresponds to a model posterior probability ≈ 0.95 (if the prior probabilities of the models in question are the same) [46], following Kass and Raftery [24]. Although in this data set the prior on *θ* did not impact model selection for detecting temporal signal, it did impact the magnitude of the Bayes factors.

**Table 2.** Log Bayes factors between isochronous and heterochronous models for each data set, separated by prior on effective population size, *θ*.

| Species; Clock Model | Exponential | Gamma | Log-normal |
| --- | --- | --- | --- |
| *Vibrio cholerae*; Strict Clock | 355.18 | 379.63 | 382.10 |
| *Vibrio cholerae*; Relaxed Clock | 208.97 | 439.63 | 219.60 |
| *Powassan virus* (POWV); Strict Clock | -80.63 | 32.67 | 50.29 |
| *Powassan virus* (POWV); Relaxed Clock | -221.94 | 18.79 | 27.23 |
| *Treponema pallidum*; Strict Clock | 105.80 | 2.17 | 1.85 |
| *Treponema pallidum*; Relaxed Clock | -34.37 | -1474.14 | -34.04 |

**Fig 1.**
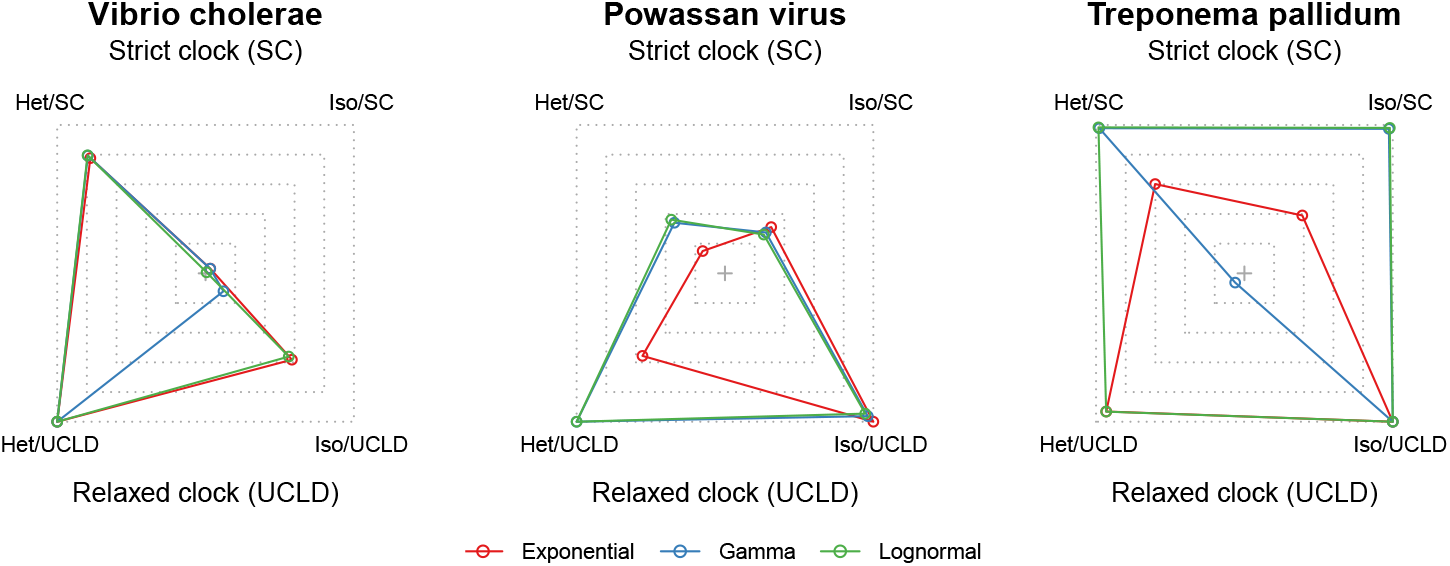
Relative log marginal likelihoods of empirical data sets. The polygons represent the relative log marginal likelihoods of each microbe data set under a different effective population size (*θ*) prior, analysed with four different configurations of sampling times and molecular clocks. The corners correspond to models (a combination of molecular clock model and the inclusion or exclusion of sampling times). The outermost dashed lines are for the highest marginal likelihood recorded and the extent to which the polygons are deformed denotes relative model support (e.g. a perfect square would imply that all models are equally well supported). A corner that falls close to the centre would correspond to a model that has low support. *Het* (heterochronous) includes sampling, while *Iso* (isochronous) does not include any sampling times. SC is strict clock and UCLD is the uncorrelated log-normal relaxed clock. Red represents an exponential prior on the effective population size, blue is a Γ prior, and green is a log-normal prior.

For our other two data sets the impact of the prior on model selection was evident. For *Poawassan virus* the Γ and log-normal priors on *θ* suggested strong temporal signal, whereas the exponential prior strongly favoured the exclusion of sampling times, according to the strict and relaxed molecular clock models. In our analyses of the *T. pallidum* data set we found support for temporal signal under the strict molecular clock, according to all priors on *θ*, although with very strong evidence only for the exponential prior and ‘positive evidence’ for the Γ and log-normal priors. Under the relaxed molecular clock model all priors had very strong support against temporal signal.

Our empirical data analyses overall demonstrate that the choice of prior may have a substantial impact on Bayesian model selection and support. We also find that the posterior distributions of key parameters, such as the evolutionary rate, were not overly sensitive to the prior for the *V. cholerae* data set (S1 Fig, *electronic supplementary material*). Moreover, this data set has been shown to have clear clocklike behaviour in root-to-tip regressions and date randomisation tests [47]. In contrast, for the data sets of *Powassan virus* and *T. pallidum*, where temporal signal was not supported for all model configurations, we found that the posterior was more sensitive to the choice of prior (S2 and S3 Figs *electronic supplementary material*).

### Simulation experiments

To understand the impact of the prior on BETS results in more detail, we conducted a set of simulation experiments where the model used to analyse the data matches the data generating process. We conducted simulations under four possible conditions: a strict or relaxed molecular clock model, and where the phylogenetic time-trees were heterochronous or isochronous. Data from heterochronous trees are sampled from a measurably evolving population and are expected to display temporal signal, whereas those from isochronous trees by definition represent a population that is no measurably evolving (the time-trees are ultrametric) and should not display temporal signal.

For data generated under heterochronous time-trees, we found that ten out of ten simulation replicates were correctly classified as having temporal signal, using a log Bayes factor of at least 3 (Table 3 and Fig 2). This perfect classification, which can be considered a low type II error (where a type II error is failing to support temporal signal when it is truly present), was supported regardless of the prior on *θ* and the molecular clock model.

**Table 3.**
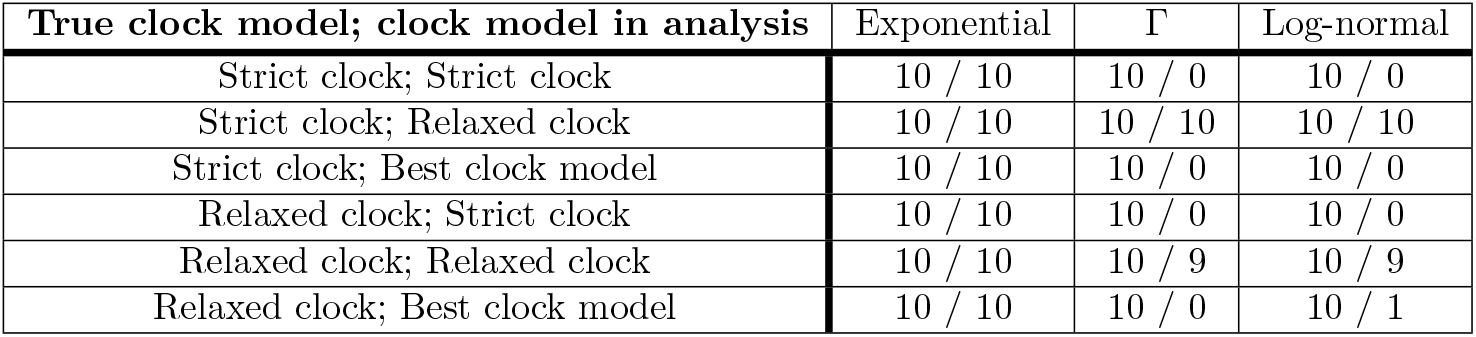
Correctly classified simulation replicates under heterochronous and isochronous trees. A total of twenty simulations were generated in each configuration. Ten under heterochronous trees that are expected to display temporal signal, and ten under isochronous trees that they are not expected to support temporal signal. A number of ten represents perfect classification according to the Bayesian evaluation of temporal signal, BETS, and a log Bayes factor of at least 3 (strong evidence for temporal signal) for heterochronous trees. For isochronous trees, the threshold is a log Bayes factor of at most -3 (strong evidence against temporal signal). The rows correspond to three possible priors on the effective population size of the constant-size coalescent, *θ*. The ‘Best clock model’ is a situation where we consider the best heterochronous and isochronous model, take their log Bayes factor, and determine temporal signal if the absolute value in favour of the correct model is at least 3. The value to the left of */* is the number of correctly classified heterochronous simulations, and the value right of */* is the number of correctly classified isochronous simulations.

| True clock model; clock model in analysis | Exponential | $\Gamma$ | Log-normal |
| --- | --- | --- | --- |
| Strict clock; Strict clock | 10 / 10 | 10 / 0 | 10 / 0 |
| Strict clock; Relaxed clock | 10 / 10 | 10 / 10 | 10 / 10 |
| Strict clock; Best clock model | 10 / 10 | 10 / 0 | 10 / 0 |
| Relaxed clock; Strict clock | 10 / 10 | 10 / 0 | 10 / 0 |
| Relaxed clock; Relaxed clock | 10 / 10 | 10 / 9 | 10 / 9 |
| Relaxed clock; Best clock model | 10 / 10 | 10 / 0 | 10 / 1 |

**Fig 2.**
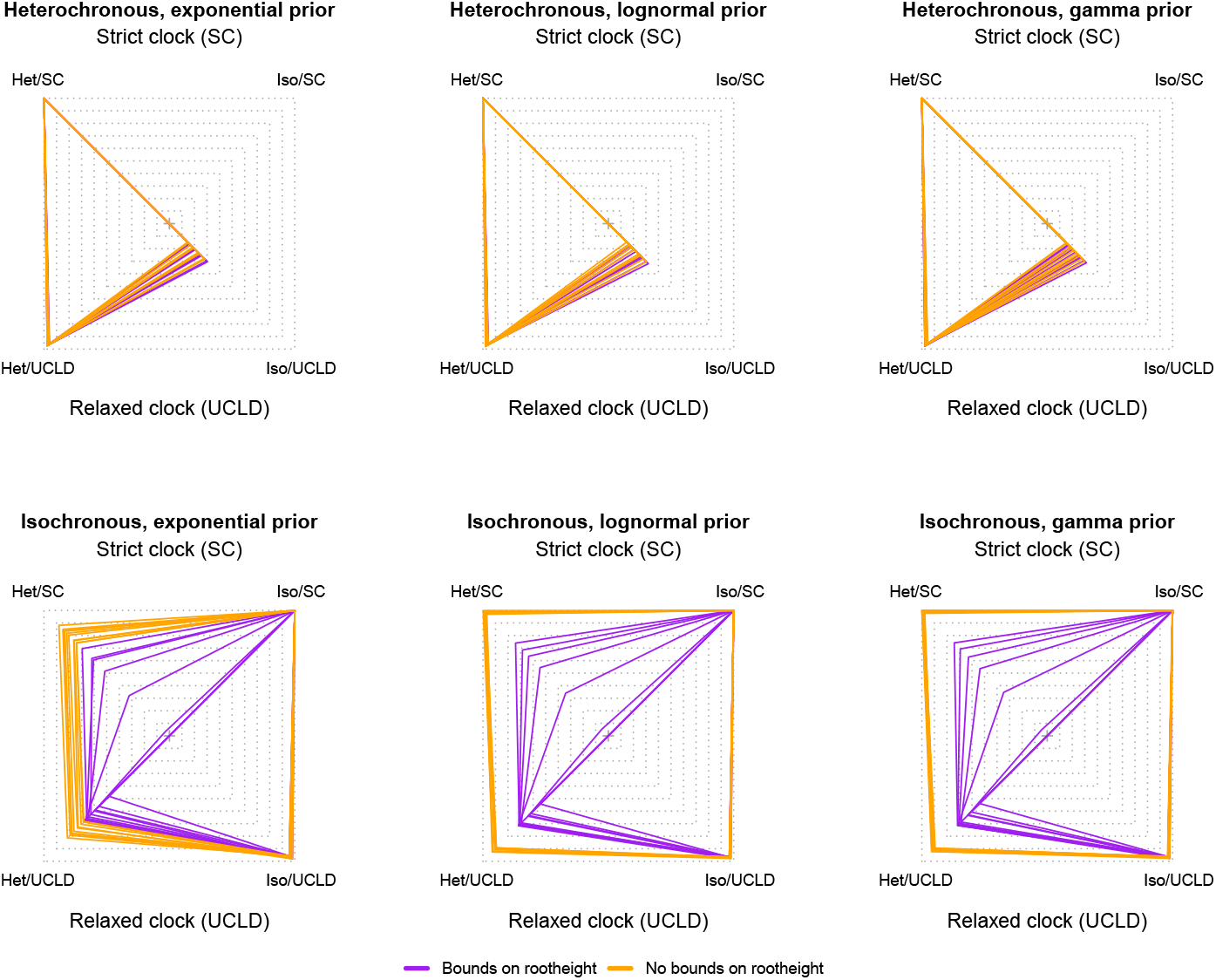
Relative log marginal likelihoods of simulations. The polygons represent the relative log marginal likelihood under three possible priors on the effective population size (*θ*) parameter of the constant-size coalescent tree prior. The top row is for heterochronous simulations, where temporal signal is present. The bottom row is for isochronous simulations that do not have temporal signal. Within each panel the corners correspond to a combination of model and sampling times, either a strict (SC) or relaxed molecular clock with an underlying log-normal distribution (UCLD), and with (heterochronous) or without (isochronous) sampling times. The correct model used to generate the data is the SC heterochronous (SC/het) for the top row and the SC isochronous (Iso/SC) for the bottom row. Each polygon is for one simulation replicate (a total of ten) and the colours denote whether we employed a hard bound on the root height of the form Uniform(0.0, 5.0), as shown in the legend.

For our data generated under isochronous trees (with no temporal signal) we found perfect classification under the exponential prior on *θ* under both clock models (Table 3 and Fig 2). For most analyses under the Γ and log-normal priors on *θ* we found that BETS incorrectly supported the presence of temporal signal, implying a high type I error (the incorrect detection of temporal signal). The exceptions were for analyses of data analysed under a relaxed molecular clock model, regardless of the molecular clock model used for simulation.

A perplexing result occurs under the best molecular clock model for the Γ and log-normal priors on *θ*. Here we take the log Bayes factor of the best heterochronous model vs the best isochronous model, which produced an increase in the classification error, relative to using the relaxed clock only. This phenomenon likely occurs because the incorrect inclusion of sampling times can mislead molecular clock model selection.

As a case in point, one of the simulation replicates under a relaxed molecular clock and an isochronous tree (with no temporal signal) had the following log marginal likelihoods; -4109.87 for the heterochronous analyses with a strict clock, -4117.06 for the isochronous analyses with a strict clock, -4124.35 for the heterochronous analyses with a relaxed clock, and -4118.29 for the isochronous analyses with a relaxed clock. The log Bayes factors under the relaxed clock have very strong evidence against temporal signal (log Bayes factor=-6.06 for heterochronous vs isochronous), whereas the opposite is true for the strict clock (log Bayes factor=7.19). However, the best heterochronous model has stronger support than the best isochronous model (here the strict or relaxed molecular clock, whose log marginal likelihoods differ by only 1.3 log likelihood units). It is also worthwhile to note that in general, analyses under the relaxed clock tended to have fewer type I classification errors (for data sets with no temporal signal) than the strict clock, regardless of the true molecular clock model used to generate the data.

Our simulation results demonstrate that detecting temporal signal when it is not present (type I error), is more common under some prior configurations (here the Γ and log-normal priors on *θ*) than the opposite (type II error, failing to detect temporal signal when it is present). Upon inspecting the resulting phylogenetic trees and the posterior of key parameters we found a probable cause. The incorrect inclusion of sampling times produces a dramatic overestimation of the height of the tree (of the root node), especially under the strict molecular clock model, a phenomenon that we refer to as ‘tree extension’ (Fig 3). Under this situation, the sampling times represent such a small proportion of the root height (the time from the most recently sampled node and the root-node), that the heterochronous tree is indistinguishable from one that is ultrametric and thus the log marginal likelihoods of a model with sampling times can be comparable, or higher than that for a model without sampling times. In fact, in Fig 3 the sampling times span 0.5 units of time, representing only 0.05% of the total height of a root height of 1,000 units of time. The phenomenon of tree extension also occurs under the relaxed molecular clock, but to a much lesser extent, and thus under this model it is easier to correctly classify isochronous data sets, because incorrect sampling times are effectively penalised. For data sets that do have temporal signal, the root height is estimated correctly, and thus tree extension does not occur Fig 4.

**Fig 3.**
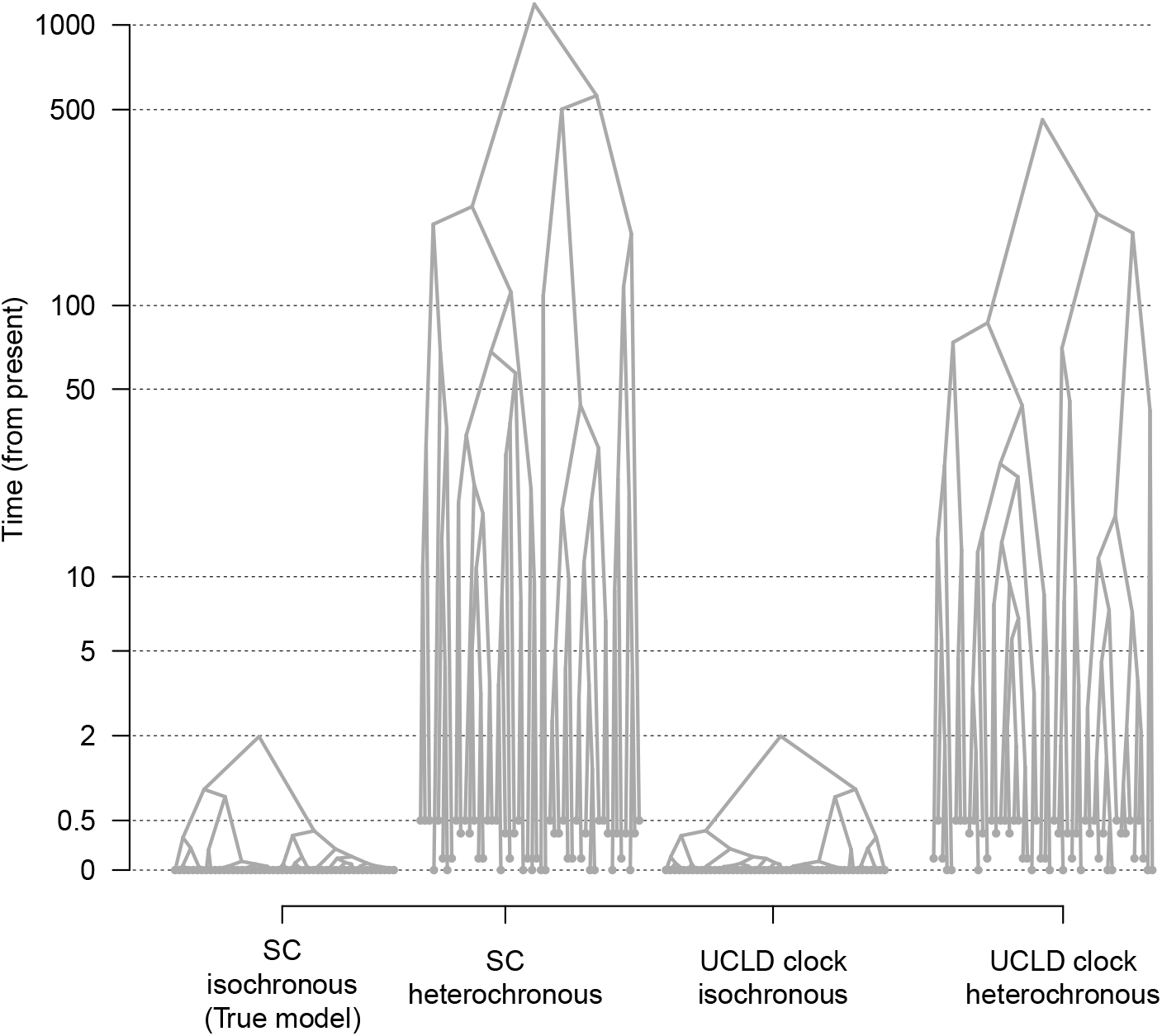
Phylogenetic tree extension for a simulation replicate with no temporal signal. Highest clade credibility trees from a data set simulated with no sampling times (isochronous) and under a strict molecular clock model (SC). The prior on effective population size (*θ*) is a Γ(*κ* = 0.001, *θ* = 1000), which resulted in high classification errors using BETS. The y-axis is the time from the present. Tip nodes have solid grey circles. Including sampling times that span 0.5 units of time and about 1/4 of the true root height induces dramatic overestimation of the root height, compared to the true model (SC, isochronous). This effect occurs under both molecular clock models, the SC and the relaxed molecular clock with an underlying log-normal distribution (UCLD), but it is markedly less pronounced in the UCLD. Note that the y-axis is in logarithmic scale (log_10_).

**Fig 4.**
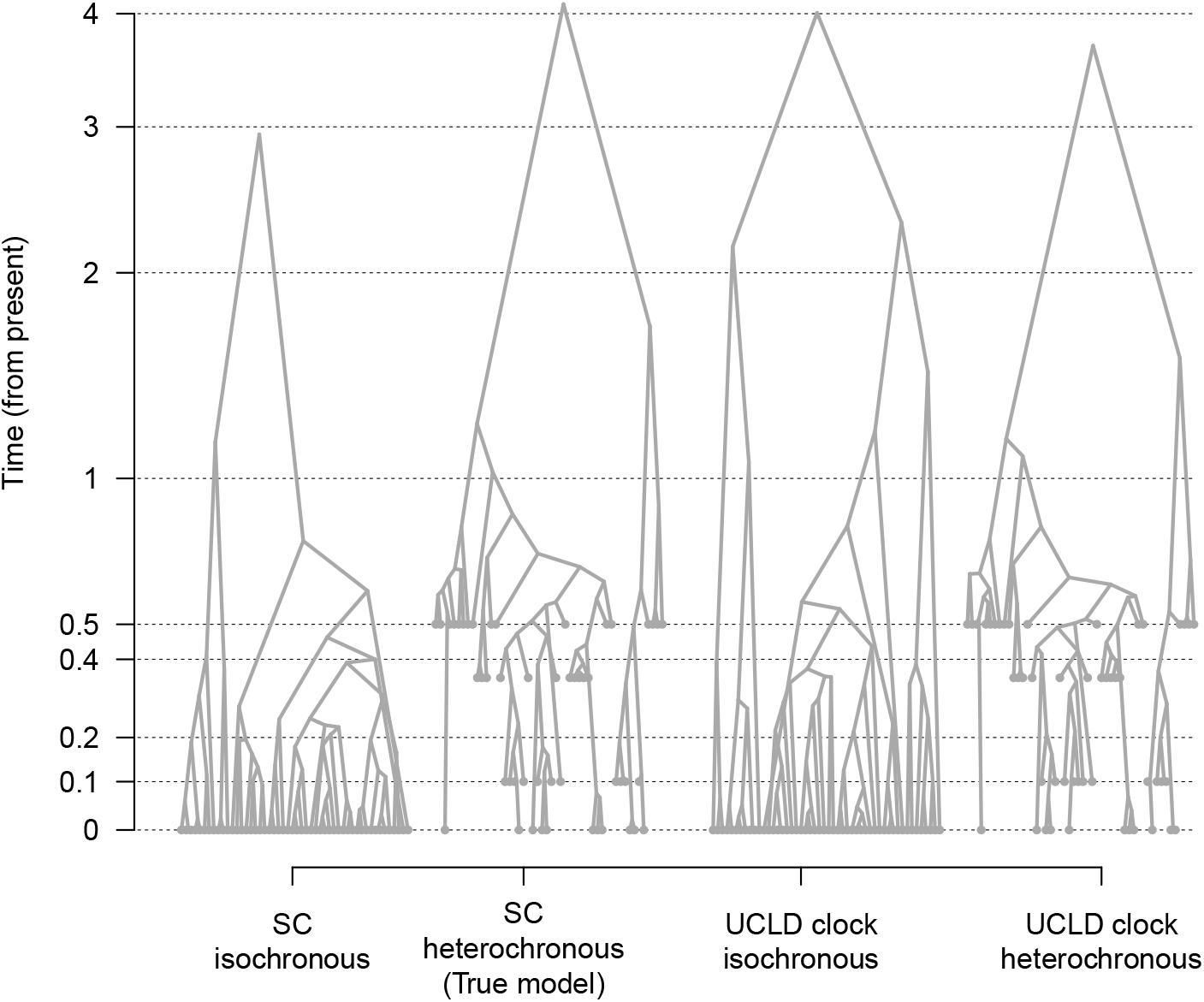
Phylogenetic trees from a simulation replicate with temporal signal. Highest clade credibility trees from a data set simulated with sampling times (heterochronous) and under a strict molecular clock model (SC). The isochronous trees are inferred by fixing the molecular clock rate to the true value, such that the timescale is in the comparable units to the heterochronous analyses. Unlike the estimates for isochronous trees (e.g. Fig 3), the height of the trees under all scenarios are comparable. Axes and labels are the same as those of Fig 3.

In isochronous analyses it is common practice to fix the clock rate to 1.0 in isochronous analyses (it is the default in BEAST1.10 and BEAST2.5 [48]), which means that the branch lengths of the time-tree will be expressed in units of subs/site. This poses a problem because the *θ* parameter of the constant-size coalescent is typically proportional to units of time [36, 49] (as are most other parameters of the tree prior, for example the growth rate of the exponential-growth coalescent). When time-calibrations are used *θ* corresponds to the population size multiplied by the generation time (*N*_*e*_ *×* generation time). Thus, the prior on *θ* for the heterochronous and isochronous analyses has different meaning (the branch lengths are not in the same units). A simple solution is to ensure that this parameter is scaled to match the units of the branch lengths, or to fix the molecular clock rate in the isochronous analyses to a biologically meaningful value. In our simulations we fixed the molecular clock rate to its true value, but we also found that using a number within the expected order of magnitude of the organism in question is sufficient (e.g. 10^−4^ to 10^−3^ subs/site/year for a ssRNA virus).

Our results indicate that using priors that favour plausible node heights is important to ensure the accuracy of temporal signal detection. However, the interplay between the parameters of the tree prior and the resulting tree topologies and node heights is not necessarily trivial, particularly when the tree prior involves multiple parameters. Thus, defining suitable priors for the parameters of the tree prior requires careful attention. A pragmatic solution is to include additional prior information in the form of hard bounds on the root height or the molecular clock rate. To this end, we investigated the effect of including a uniform prior between 0.0 and 5.0 units of time for the root height, meaning that trees that are older than 5.0 units have a prior probability of 0.0. Crucially, the trees under which we generated our simulated data had root heights of around 2.0 units of time, and thus the hard bound of 5.0 allows for trees that are over twice as old as the truth.

Setting hard bounds on the root height resulted in perfect classification accuracy for both the heterochronous and isochronous simulations (Table 4 and Fig 2, respectively). The improvement in classification for data sets with no temporal signal is likely because the hard bounds prevent the tree extension phenomenon. This allows the inclusion of sampling times to impose a penalty on the log marginal likelihood (see polygons in Fig 2). Our empirical data set of *V. cholerae*, which had evidence of temporal signal under all prior conditions, consistent with previous analyses [30], also displayed strong evidence for temporal signal with a hard bound on of 500 years before present on the root height (all log Bayes factors were at least 200 in favour of temporal signal; S4 Fig, *electronic supplementary material*).

**Table 4.** Correctly classified simulation replicates under heterochronous and isochronous trees using hard bounds on the root height. Rows and columns are identical to those of Table 3, but here the analyses include an explicit prior on the root height, via a uniform distribution between 0 and 5.0.

| True clock model; clock model in analysis | Exponential | $\Gamma$ | Log-normal |
| --- | --- | --- | --- |
| Strict clock; Strict clock | 10 / 10 | 10 / 10 | 10 / 10 |
| Strict clock; Relaxed clock | 10 / 10 | 10 / 10 | 10 / 10 |
| Strict clock; Best clock model | 10 / 10 | 10 / 10 | 10 / 10 |
| Relaxed clock; Strict clock | 10 / 10 | 10 / 10 | 10 / 10 |
| Relaxed clock; Relaxed clock | 10 / 10 | 10 / 10 | 10 / 10 |
| Relaxed clock; Best clock model | 10 / 10 | 10 / 10 | 10 / 10 |

The exponential-growth coalescent is more flexible than the constant-size coalescent, where population size change is allowed to change deterministically (for its description in a phylodynamic context see [50, 51]). This model has two key parameters, the ‘scaled population size’ (Φ) and the exponential growth rate. Due to its coalescent nature the time to the most recent common ancestor scales positively with Φ. We investigated the performance of BETS under this tree prior, using the same three prior distributions on Φ as we did for *θ* in the constant-size coalescent and for the growth rate we used *Laplace*(*µ* = 0.0, *b* = 1.0). The relative log marginal likelihoods between models were very similar to those using the constant-size coalescent. Heterochronous data sets were overwhelmingly classified as having temporal signal, whereas isochronous data sets required hard bounds to correctly detect a lack of temporal signal (S5 Fig, *electronic supplementary material*).

To investigate the phenomenon of tree extension in more realistic conditions, we conducted ten simulations of data that resembled our *V. cholerae* empirical data (similar numbers of variable sites and distribution of sampling times), and evaluated them using BETS under an exponential prior on *θ*. We only considered isochronous trees under a strict clock, because this is the situation where we obtained the largest number of false positives (type I errors). In the absence of hard bounds on the root height five out of ten data sets resulted in false positives (‘strong’ support for temporal signal), with those that were misclassified displaying tree extension. The use of hard bounds (five fold older than the true root height) reduced the number of false positives to one.

### Prior predictive simulations and parameter correlations

Our finding that the tree prior can affect model selection prompted an investigation of parameter interactions and of the expectations under the prior. We simulated phylogenetic trees from a prior distribution to inspect the correlation of parameters and the marginal prior for those that have obvious associations (e.g. the evolutionary rate has a CTMC-rate reference prior that depends on tree length). These simulations are commonly known as prior predictive simulations (e.g. [52]), and referred to as ‘sampling from the prior’ in the phylogenetics literature [53]. Initially, we set a uniform prior on *θ* from 0 to 10^3^ and recorded the evolutionary rate, the tree length and root height. This prior is not generally recommended [54], and we present it here to illustrate correlations between parameters and the marginal prior, instead of using it for analysing our data.

An obvious finding is a natural positive correlation between tree length and root height, although the trend is not strictly linear (Fig 5). We also observed a positive correlation between *θ*, tree length and root height, which is also expected because large population sizes impose long coalescent times [55]. The nature of this trend is heteroskedastic, with the variance in these tree statistics increasing with *θ*. Our simulations demonstrate an inverse relationship between the evolutionary rate, *θ* and the tree statistics, but with a range that can span several orders of magnitude and with values ranging from 10^−9^ to 10^−4^ subs/site/time, meaning that the CTMC-rate reference prior tends to be diffuse. It is also noteworthy that the uniform prior on *θ* does not result in uniform marginal prior distributions for any of the parameters investigated here. These results illustrate the importance of visualising the prior and the fact that it is difficult to predict how different parameters will interact.

**Fig 5.**
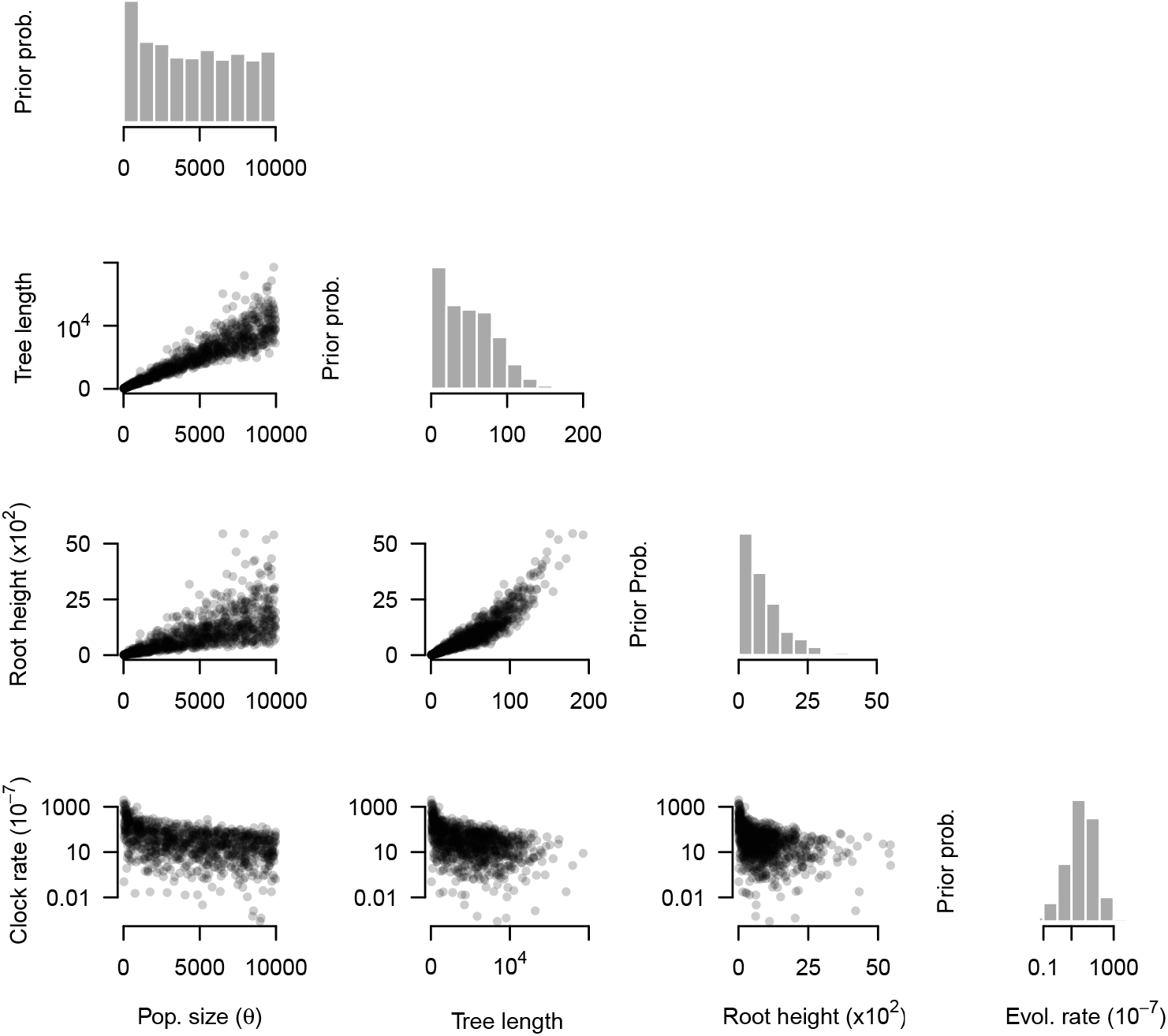
Marginal prior distributions and pairs plots. The grey histograms correspond to the parameter labelled at the bottom of each column; effective population size (pop. size, *θ*), tree length, root height, and the evolutionary rate (evol. rate). The prior for *θ* here is a Uniform distribution between 0 and 10^3^, while that for the evolutionary rate is a CTMC-rate reference prior. Note that the tree length and root height have units of time, the evolutionary rate is in subs/site/time, and *θ* is proportional to units of time.

We conducted prior predictive simulations for the six prior configurations for *θ* under a heterochronous data set. In Fig 6 we show the resulting distribution on the root height. The exponential (*θ* ∼ *Exponential*(*µ* = 1.0), Γ (Γ(*κ* = 0.001, *θ* = 1000)) and log-normal (log-normal(*µ* = 1.0, *σ* = 5.0)) prior distributions respectively produced mean root heights of 2.90 (95% quantile range, qr: 1.62 to 14.34), 1.61 (95% qr: 1.50 to 4.87) and 772.11 (95% qr: 1.66 to 5.06 *×* 10^5^) units of time. Clearly, the log-normal is the most vague prior here, but it produces implausible values of several orders of magnitude higher than our expectation of root heights of around one and ten units of time (as simulated using *θ* = 1.0).

**Fig 6.**
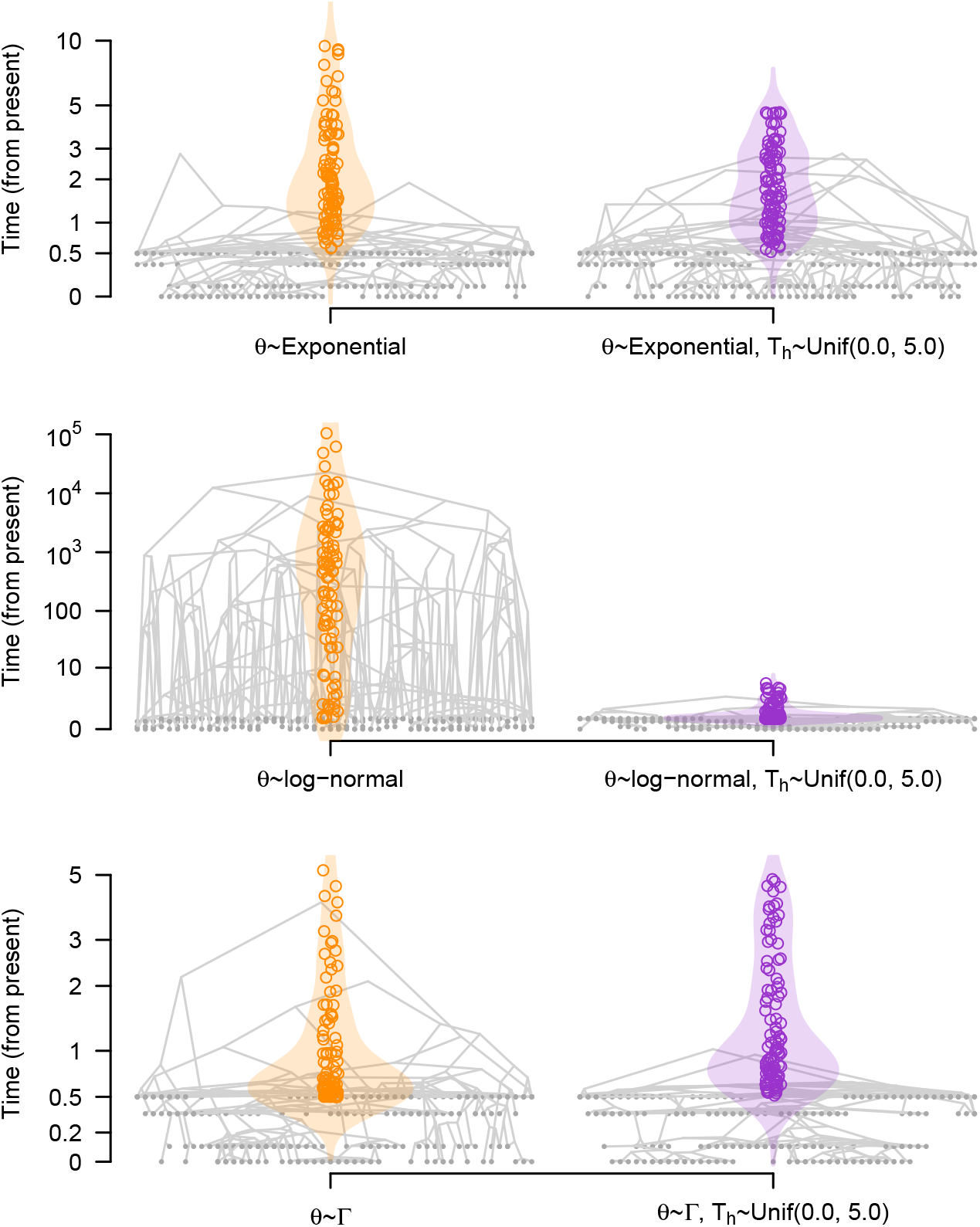
Prior predictive simulations and marginal priors of root heights, given the prior on the effective population size, *θ*. Each panel corresponds to a different prior on *θ*, as described on Table 1 (Exponential(*µ* = 1.0), Log-normal(*µ* =1.0, *σ* =5.0), and Γ(*κ* = 0.001, *θ* = 1000)). We show five simulated trees from our analysis using sampling times (heterochronous analyses) and overlaid them (similar to densitree plots [56]). The violin plots show the prior densities of the root height and the hollow circles denote 100 randomly drawn samples from the prior. The y-axis is the time from the present, but note that the scales are different. Tip nodes are shown with solid grey circles. The densities and trees on the left, in orange, do not include an explicit prior on the root height (*T*_*h*_), while those to the right, in purple, have a hard bound on the root height in the form of a uniform prior between 0.0 and 5.0 units of time.

For comparison, we also simulated trees under the same priors on *θ*, with hard bounds on the root height, and with a uniform distribution with minimum and maximum values of 0.0 and 5.0, as described above. In this case, the exponential prior on *θ* yielded trees with root heights of mean 2.47 (95% qr: 1.62 to 5.16), whereas those for the log-normal and Γ were 1.61 (95% qr: 1.50 to 5.11), and 1.96 (95% qr: 1.54 to 4.42), respectively. As a result, in the log-normal prior on *θ* the use of hard bounds on the root height resulted in much shorter root heights, with a smaller impact on the exponential and Γ priors.

## Discussion

Our study demonstrates that the choice of prior distribution is a key factor for Bayesian model selection and that it can mislead tests of temporal signal. In general, data sets with no temporal signal are easily misclassified when the prior favours an implausibly old root height and low evolutionary rates, resulting in type I errors. The incorrect detection of temporal signal may lead to a systematic overestimation of the evolutionary timescale and an underestimation of molecular clock rates.

We find that tree extension is the most probable reason for type I errors in BETS, because it reduces the sampling window relative to the root height, and therefore trees with sampling times have phylogenetic likelihoods that are very similar to those of ultrametric trees (S6 Fig, *electronic supplementary material*). Because the phylogenetic likelihood has substantial weight on the marginal likelihood, this phenomenon can mislead model selection. Verifying that the prior does not favour trees that are implausibly old can mitigate this problem. In our simulation study, the exponential prior on *θ* reduced the degree of tree extension, but this particular prior might not be reasonable for some data sets, such as those where we expect very old divergences between sequences or large population sizes.

Our observation of tree extension pertains to isochronous data sets analysed with sampling times, but a similar situation occurs in date-randomisation tests [18]. Here, the sequence sampling times are permuted a number of times, the evolutionary rate is re-estimated, and the data are considered to have temporal signal if the estimates from the permutations do not overlap with that from the correct sampling times. This test exists in a Bayesian and maximum-likelihood setting, where point estimates are used instead of posterior distribution [21]. Notably, when the data do have temporal signal, the estimates from the permutations are substantially lower, implying older times to the most recent common ancestor [17]. Therefore the inclusion of incorrect sampling times, whether the data set is truly isochronous or not, may result in tree extension as a means of compensating for the likelihood penalty imposed by incorrect sampling times.

While the phenomenon of tree extension occurs for the strict and relaxed molecular clock models, it is less pronounced in relaxed molecular clock models (e.g. see Fig 3). We hypothesise that the relaxed molecular clock is more robust to tree extension in data sets that are truly isochronous because this model can absorb the incorrect inclusion of sampling times by treating them as rate variation among lineages. However, careful molecular clock model selection is advisable for tests of temporal signal and molecular dating in general.

The presence, rather than the absence, of temporal signal is much easier to detect, meaning low type II errors in BETS. In this case, specifying an incorrect isochronous model does not result in an obvious distortion of the time tree that could mislead BETS, at least under the conditions that we used here. It is conceivable, however, that certain priors on the phylogenetic tree or evolutionary rate could result in type II errors.

A key consideration is that the parameters of the tree prior for isochronous and heterochronous analyses should be in the same units (and of similar magnitude), which facilitates comparison of the models in question. For example, fixing the evolutionary rate to 1.0 in an isochronous analysis means that branch lengths of the time tree are in units of expected genetic distance (usually subs/site) and thus the coalescent parameters have a different meaning to those estimated under heterochronous analyses (where the evolutionary rate is a free parameter). A more tractable approach is to conduct isochronous analyses by fixing the evolutionary rate to a value within the expected order of magnitude, as we have done here, or rescaling the parameters of the tree prior, such as *θ*, accordingly (in units that are the proportional to time^−1^; see [35]). Biological knowledge, such as the negative correlation between genome size and rates of microbes [47, 57] or simply a previous estimate for a closely related organism, can be helpful for specifying pausible values for the molecular clock rate.

Our analyses only considered fully parametric coalescent models for the tree prior. This is a convenient choice for the purpose of our experiments, but molecular clock dating can be performed using other tree priors, which are amenable to using BETS, under the condition that the prior is proper. For example, current implementations of the Bayesian skyline plot do not constitute a proper prior, unlike the Bayesian skyride that is fully proper (for an in-depth discussion see [58]). Although birth-death tree priors could conceivably be used in the context of BETS (they are usually fully parametric and have proper prior distributions), these models explicitly use sampling time information as data [59]. Clearly, more work is needed to determine whether comparing the log marginal likelihoods of the isochronous and heterochronous versions of birth-death models is valid.

Our results point to a few recommendations to improve tests of temporal signal. First, researchers should carefully elicit the priors, which appears important for Bayesian model selection (see [60] for a related problem in tree topology tests). Prior predictive simulations have been recommended for assessing interactions between multiple calibration priors [61]. We find that they are also essential to understand potential interactions between the different parameters in the model and ultimately whether the prior expectation is reasonable with respect to our knowledge about the data. The log-normal and exponential priors on *θ* produced trees with root heights that differed by over four orders of magnitude. In our simulation study, using hard bounds on the root height alleviated the problem of tree extension when the prior on *θ* favoured very old trees. In empirical data, however, one may not want to make such strong statements, and prior predictive simulations can help determine whether the tree prior and associated parameters produce trees with sensible root heights (e.g. the exponential prior on *θ* had low type I and type II errors in our simulations). As a case in point, for a data set collected from a recently emerging pathogen the exponential prior on *θ* will likely generate more reasonable trees than the log-normal. Conversely, the log-normal prior, as parameterised here, might generate more reasonable trees for a data set involving ancient DNA samples.

Our second recommendation is to conduct prior sensitivity analyses on a set of candidate priors to determine the extent to which the prior can influence the posterior [62, 63], and model selection using Bayes factors [64]. Visualising the posterior and prior distributions for a range of priors can be illuminating (see S1 through S3 Figs, *electronic supplementary material*). A prior that makes unreasonable statements and is overly influential on the posterior may need to be revised and a comparable approach has been proposed for phylogeographic models [42, 43]. The exponential prior on *θ* in the *Powassan virus* did not support temporal signal, but note that it is much more influential on the posterior than the Γ and log-normal priors for this particular data set. In contrast, the *V. cholerae* data seemed robust to the three priors that we used. Some empirical data sets yielded unclear evidence for temporal signal according to BETS, such as the *T. pallidium* data set that we reanalysed. In such cases, the decision of whether the data have sufficient temporal signal may require multiple lines of evidence, such as date-randomisation tests [18, 19], comparisons of the prior and posterior [65], and root-to-tip regressions. [13, 14]. When temporal signal is inconclusive, the inclusion of additional information, such as internal node calibrations and informative molecular clock rate priors may be essential for molecular dating - as was the case in that study [31]. Our use of hard bounds on the root height is effectively an internal node constraint, but a similar calibration priors in the form of a parametric distributions on any other internal node or nodes could be used [61, 66].

Finally, we find that the choice of the molecular clock model has a tangible impact on tests of temporal signal. The strict molecular clock model is more prone to type I error than the relaxed molecular clock model. The most likely reason for this finding is that this approach mitigates tree extension when the data have no temporal signal. Thus we recommend that the molecular clock be selected carefully.

The sum of our results and recommendations has implications for the reliability of estimates of evolutionary rates and timescales, for determining whether a population is measurably evolving, and even for assessing the phylodynamic threshold of emerging microbes [67]. Concretely, genome data from recently emerging microbes may have very few substitutions, in which case prior sensitivity analyses and prior predictive simulations would easily reveal whether the data are sufficiently informative. Overall, tests of temporal signal have a key place in genomic analyses of pathogens and our results will be useful as guidelines to improve their application and interpretation.

## Materials and methods

### Empirical data

We selected three different data sets to evaluate temporal signal, *V. cholerae* [30], *Powassan virus* [32], and *T. pallidum* [31]. We chose these data sets because they included ancient samples (*T. pallidum* and *V. cholerae*) and very divergent sequences (*Powassan virus*), such that they have the complexity of many real-world analyses. Our data and example code for our analyses is available at https://github.com/sebastianduchene/BETS_code_data.

The sequence alignments varied in their sampling timeframe, number of informative sites, and number of sequences. The most sparse data set was *T. pallidum*, where sampling dates spanned 481 years (1534 to 2016), with 28 sequences each containing 1,500 SNP sites. Among these samples, there were only two in the 1500’s and two in the 1700’s, with the rest of the samples concentrated in the 1900’s to 2000’s. There were 319 sequences with 11,193 sites (the complete genome) for *Powassan virus*, spanning 24 years of sampling times (1995 to 2019). Most of the samples were collected after 2010, with only three isolated in 1995. For *V. cholera* we had 122 sequences with 1,57 SNP sites across 73 years of sampling (1937 to 2010). There are 1,392 unique site patterns in *cholera*, 3,457 in *Powassan virus*, and 840 in *T. pallidum*.

For *V. cholerae* and *T. pallidum* our data consisted of SNP sites. To account for such ascertainment bias, we specified the number of constant nucleotides in the whole genome in our subsequent analyses. We sampled the posterior distribution using Markov chain Monte Carlo as implemented in BEAST1.10. The chain length was 10^8^ steps, sampling every 10^3^ to draw a total of 10^4^ samples. The prior for the evolutionary rate of the strict clock was a CTMC-rate reference prior (i.e. a Γ(*α* = 0.5, *β* =tree length), and mean=*α/β*) [41]. For the log-normal distribution of the relaxed clock we also used the CTMC-rate reference prior for the mean rate (known as the ucld.mean in the program) and an exponential prior with mean 0.33 for the standard deviation (the ucld.stdev in the program). For the tree prior we used a constant-size coalescent with population size, *θ* with three possible priors, as described in Table 1. In all cases we used the HKY+Γ_4_ substitution model, with the default priors in BEAST1.10.

To calculate log marginal likelihoods we used generalised stepping-stone [68, 69]. This method requires a working distribution for all parameters, including the tree prior, for which we used the matching coalescent model. We set 100 path steps between the unnormalised posterior and the working distribution, following equally spaced intervals from a *β*(0.3, 1.0) distribution. For each step we ran a chain length of 2 *×* 10^6^ steps. We considered these settings to be appropriate after repeating the log marginal likelihood calculations 10 times for two of data sets and ensuring that the values did not vary by more than 1.0 log likelihood units.

### Simulation experiments

For our simulations we considered well-understood conditions to isolate the impact of the tree prior on tests of temporal signal. We simulated phylogenetic trees under a constant-size coalescent model with a fixed population size of 1.0 and a resulting average root height of 2.0 units of time. The trees could be isochronous (i.e. ultrametric) or heterochronous. For the latter we assigned tip heights of of 0.0, 0.10, 0.35, or 0.50, with equal probability. Our simulations simplify the complexity or empirical data (e.g. simple substitution model, no population structure), but they represent a realistic situation where data were collected across four discrete sampling periods, which resembles sequencing blitzes [70], or archaeological sampling of strata [71]. Because the coalescent model used to analyse the data is conditioned on sampling times we expect it to be robust to such sampling bias [39, 72, 73]. We used a JC substitution model [74] (all exchangeability parameters are equal) to generate sequence alignments of 1,000 nucleotides and an evolutionary rate of 0.05 subs/site/time, which resulted in around 250 unique site patterns. For our simulations under a relaxed molecular clock model we sampled branch rates from a log-normal distribution with mean 0.05 and standard deviation of 0.25. The procedure for obtaining the simulated alignments consisted of specifying the model above in BEAST1.10 with an empty sequence alignment to sample phylogenetic trees. We then used NELSI [75] and Phangorn [76] to simulate evolutionary rates (a single value for the strict clock and the branch rates for the relaxed molecular clock model) and sequence alignments, respectively.

We analysed the simulated data under heterochronous and isochronous models. For the heterochronous analyses we used the correct sampling times (where the correct model was the heterochronous) or assigned the tip heights above randomly (0.0, 0.10, 0.35, or 0.50) where the isochronous was the true model. For the evolutionary rate (clock.rate in the strict and ucld.mean for the relaxed molecular clock model) we set the default CTMC-rate reference prior and the six possible configurations of the prior on *θ* (Table 1, plus those with hard bounds on the root height). For the isochronous analyses we did not specify sampling times and we fixed evolutionary rate to its true value of 0.05 to ensure that the branch lengths, *θ* and other parameters are in the correct units. We calculated log marginal likelihoods with the same procedure as for the empirical data.

We also generated a set of simulations under conditions aimed at mimicking the empirical *V. cholerae* data set. We focused our attention on isochronous simulations, which resulted in the highest type I errors in all other conditions. We simulated phylogenetic trees with 50 taxa under a coalescent process with a constant population size, *θ*, of 100.0. The root height of these trees ranged between 60.0 and 248.0 units of time. To specify an artificial set of sampling times we sampled from an exponential distribution with mean of 1/10^th^ of the root height. This approach resulted in sampling times that spanned about 1/3^rd^ of the height of the tree, with most samples collected close to the present. We set the clock rate and genome length to produce a similar number of site patterns as in the empirical data [30]. We conducted our analyses using the same methods as the rest of the simulations above, except that in this case we only used the exponential prior on *θ* with a mean of 100.0 for *θ* (the value under which we generated the data), and with and without hard bounds on the root height.

## Supporting information

**S1 Fig. Densities of key statistics of the *Vibrio cholerae* empirical data**. The clock rate, root height, tree length, and coefficient of rate variation are shown under three priors on *θ*, exponential (red), gamma (blue), and log-normal (green).

**S2 Fig. Densities of key statistics of *Powassan virus* empirical data**. The clock rate, root height, tree length, and coefficient of rate variation are shown under three priors on *θ*, exponential (red), gamma (blue), and log-normal (green).

**S3 Fig. Densities of key statistics of *Treponema palladium* empirical data**. The clock rate, root height, tree length, and coefficient of rate variation are shown under three priors on *θ*, exponential (red), gamma (blue), and log-normal (green).

**S4 Fig. Relative log marginal likelihoods of empirical data sets with bounds on root height**. The polygons represent the relative log marginal likelihoods of each microbe data set under a different effective population size (*θ*) prior, analysed with four different configurations. *Het* (heterochronous) includes sampling, while *Iso* (isochronous) does not include any sampling times. SC is strict clock and UCLD is the uncorrelated log-normal relaxed clock. Red represents an exponential prior on the effective population size, blue is a Γ prior, and green is a log-normal prior.

**S5 Fig. Relative log marginal likelihoods of simulations analysed under an exponential-growth coalescent tree prior**. The top row is for heterochronous simulations, where temporal signal is present, and the bottom row is for isochronous simulations that do not have temporal signal. Within each panel the corners correspond to a combination of model and sampling times, either a strict (SC) or relaxed molecular clock with an underlying log-normal distribution (UCLD), and with (heterochronous) or without (isochronous) sampling times. The polygons represent the relative log marginal likelihood under three possible priors on the scaled population size (Φ) parameter of the exponential-growth coalescent tree prior. The correct model used to generate the data is the SC heterochronous (SC/het) for the top row and the SC isochronous (Iso/SC) for the bottom row. Each polygon is for one simulation replicate (a total of ten) and the colours denote whether we employed a hard bound on the root height of the form Uniform(0.0, 5.0), as shown in the legend.

**S6 Fig. The impact of tree extension in the phylogenetic likelihood**. Phylogenetic likelihood (i.e. the probability of the sequence data, given the full model and the prior) vs the root height of phylogenetic trees. Each panel represents a data set that was truly isochronous and the prior on the effective population size of the constant-size coalescent, *θ*. Here we show the posterior distributions for a single data set, analysed under three different priors on *θ* and with sampling times that were isochronous (no sampling times), or heterochronous, and with and without hard bounds on the root height. Each point corresponds to a sample from the posterior, with shapes and colours described in the legend. Note that the isochronous (orange circles) and heterochronous without bounds have similar phylogenetic likelihoods, despite having trees with root heights that differ by multiple orders of magnitude. In practice, the heterochronous analyses without bounds produce trees with root heights that are so old that they are effectively isochronous (see Fig 6). In contrast, using hard bounds on the root height, even when the bounds are twice the true age of the root height results in a substantial penalty on the phylogenetic likelihood, and the correct classification in BETS. Note that here the exponential prior on *θ* results in much younger root height values than for the two other priors on this parameter.

## References

1. Bromham L, Duchêne S, Hua X, Ritchie AM, Duchêne DA, Ho SY. Bayesian molecular dating: opening up the black box. Biological Reviews. 2018;93(2):1165–1191.

2. Hong Q, Chen G, Tang ZZ. PhyloMed: a phylogeny-based test of mediation effect in microbiome. Genome Biology. 2023;24(1):72.

3. Zhou C, Zhao H, Wang T. Transformation and differential abundance analysis of microbiome data incorporating phylogeny. Bioinformatics. 2021;37(24):4652–4660.

4. Zuckerkandl E, Pauling L. Evolutionary divergence and convergence in proteins. In: Evolving genes and proteins. Elsevier; 1965. p. 97–166.

5. Drummond AJ, Ho SYW, Phillips MJ, Rambaut A. Relaxed phylogenetics and dating with confidence. PLoS Biology. 2006;4(5):e88.

6. Ho SY, Duchêne S. Molecular-clock methods for estimating evolutionary rates and timescales. Molecular Ecology. 2014;23(24):5947–5965.

7. Yang Z, Rannala B. Bayesian estimation of species divergence times under a molecular clock using multiple fossil calibrations with soft bounds. Molecular biology and evolution. 2006;23(1):212–226.

8. Dos Reis M, Yang Z. The unbearable uncertainty of Bayesian divergence time estimation. Journal of Systematics and Evolution. 2013;51(1):30–43.

9. Rodrigo AG, Felsenstein J. Coalescent approaches to HIV population genetics. The evolution of HIV. 1999; p. 233–272.

10. Korber B, Muldoon M, Theiler J, Gao F, Gupta R, Lapedes A, et al. Timing the ancestor of the HIV-1 pandemic strains. science. 2000;288(5472):1789–1796.

11. Drummond AJ, Pybus OG, Rambaut A, Forsberg R, Rodrigo AG. Measurably evolving populations. Trends in ecology & evolution. 2003;18(9):481–488.

12. Biek R, Pybus OG, Lloyd-Smith JO, Didelot X. Measurably evolving pathogens in the genomic era. Trends in ecology & evolution. 2015;30(6):306–313.

13. Rambaut A, Lam TT, Max Carvalho L, Pybus OG. Exploring the temporal structure of heterochronous sequences using TempEst (formerly Path-O-Gen). Virus evolution. 2016;2(1):vew007.

14. Featherstone LA, Rambaut A, Duchene S, Wirth W. Clockor2: Inferring Global and Local Strict Molecular Clocks Using Root-to-Tip Regression. Systematic Biology. 2024; p. syae003. doi:10.1093/sysbio/syae003.

15. Volz E, Frost SD. Scalable relaxed clock phylogenetic dating. Virus evolution. 2017;3(2):vex025.

16. Doizy A, Prin A, Cornu G, Chiroleu F, Rieux A. Phylostems: a new graphical tool to investigate temporal signal of heterochronous sequences datasets. Bioinformatics Advances. 2023;3(1):vbad026.

17. Rieux A, Balloux F. Inferences from tip-calibrated phylogenies: a review and a practical guide. Molecular ecology. 2016;25(9):1911–1924.

18. Ramsden C, Holmes EC, Charleston MA. Hantavirus evolution in relation to its rodent and insectivore hosts: no evidence for codivergence. Molecular biology and evolution. 2009;26(1):143–153.

19. Duchêne S, Duchêne D, Holmes EC, Ho SY. The performance of the date-randomization test in phylogenetic analyses of time-structured virus data. Molecular Biology and Evolution. 2015;32(7):1895–1906.

20. Murray GG, Wang F, Harrison EM, Paterson GK, Mather AE, Harris SR, et al. The effect of genetic structure on molecular dating and tests for temporal signal. Methods in Ecology and Evolution. 2016;7(1):80–89.

21. Duchene S, Duchene DA, Geoghegan JL, Dyson ZA, Hawkey J, Holt KE. Inferring demographic parameters in bacterial genomic data using Bayesian and hybrid phylogenetic methods. BMC evolutionary biology. 2018;18:1–11.

22. Duchene S, Lemey P, Stadler T, Ho SY, Duchene DA, Dhanasekaran V, et al. Bayesian evaluation of temporal signal in measurably evolving populations. Molecular Biology and Evolution. 2020;37(11):3363–3379.

23. Gavryushkin A, Drummond AJ. The space of ultrametric phylogenetic trees. Journal of theoretical biology. 2016;403:197–208.

24. Kass RE, Raftery AE. Bayes factors. Journal of the American Statistical Association. 1995;90(430):773–795.

25. Molak M, Suchard MA, Ho SY, Beilman DW, Shapiro B. Empirical calibrated radiocarbon sampler: a tool for incorporating radiocarbon-date and calibration error into B ayesian phylogenetic analyses of ancient DNA. Molecular ecology resources. 2015;15(1):81–86.

26. Spyrou MA, Bos KI, Herbig A, Krause J. Ancient pathogen genomics as an emerging tool for infectious disease research. Nature Reviews Genetics. 2019;20(6):323–340.

27. Duchêne S, Ho SY, Carmichael AG, Holmes EC, Poinar H. The recovery, interpretation and use of ancient pathogen genomes. Current Biology. 2020;30(19):R1215–R1231.

28. Gelman A, Rubin DB. Avoiding model selection in Bayesian social research. Sociological methodology. 1995;25:165–173.

29. Gelman A, Carlin JB, Stern HS, Rubin DB. Bayesian data analysis; 2014.

30. Devault AM, Golding GB, Waglechner N, Enk JM, Kuch M, Tien JH, et al. Second-pandemic strain of Vibrio cholerae from the Philadelphia cholera outbreak of 1849. New England Journal of Medicine. 2014;370(4):334–340.

31. Majander K, Pfrengle S, Kocher A, Neukamm J, Du Plessis L, Pla-Díaz M, et al. Ancient bacterial genomes reveal a high diversity of Treponema pallidum strains in early modern Europe. Current Biology. 2020;30(19):3788–3803.

32. Vogels CB, Brackney DE, Dupuis AP, Robich RM, Fauver JR, Brito AF, et al. Phylogeographic reconstruction of the emergence and spread of Powassan virus in the northeastern United States. Proceedings of the National Academy of Sciences. 2023;120(16):e2218012120.

33. Suchard MA, Lemey P, Baele G, Ayres DL, Drummond AJ, Rambaut A. Bayesian phylogenetic and phylodynamic data integration using BEAST 1.10. Virus Evolution. 2018;4(1):vey016.

34. Heath TA, Moore BR. Bayesian inference of species divergence times. Bayesian phylogenetics: methods, algorithms, and applications. 2014; p. 277–318.

35. Boskova V, Bonhoeffer S, Stadler T. Inference of epidemiological dynamics based on simulated phylogenies using birth-death and coalescent models. PLoS Computational Biology. 2014;10(11):e1003913.

36. Drummond AJ, Nicholls GK, Rodrigo AG, Solomon W. Estimating mutation parameters, population history and genealogy simultaneously from temporally spaced sequence data. Genetics. 2002;161(3):1307–1320.

37. R Oaks J, A Cobb K, N Minin V, D Leaché A. Marginal likelihoods in phylogenetics: a review of methods and applications. Systematic Biology. 2019;68(5):681–697.

38. Baele G, Li WLS, Drummond AJ, Suchard MA, Lemey P. Accurate model selection of relaxed molecular clocks in Bayesian phylogenetics. Molecular Biology and Evolution. 2013;30(2):239–243. doi:10.1093/molbev/mss243.

39. Featherstone LA, Di Giallonardo F, Holmes EC, Vaughan TG, Duchêne S. Infectious disease phylodynamics with occurrence data. Methods in Ecology and Evolution. 2021;12(8):1498–1507.

40. Wang Y, Yang Z. Priors in Bayesian phylogenetics. Bayesian phylogenetics: methods, algorithms, and applications. 2014; p. 5–24.

41. Ferreira MA, Suchard MA. Bayesian analysis of elapsed times in continuous-time Markov chains. Canadian Journal of Statistics. 2008;36(3):355–368.

42. Gao J, May MR, Rannala B, Moore BR. Model misspecification misleads inference of the spatial dynamics of disease outbreaks. Proceedings of the National Academy of Sciences. 2023;120(11):e2213913120.

43. Gao J, May MR, Rannala B, Moore BR. PrioriTree: a utility for improving phylodynamic analyses in BEAST. Bioinformatics. 2023;39(1):btac849.

44. Höhna S, Heath TA, Boussau B, Landis MJ, Ronquist F, Huelsenbeck JP. Probabilistic graphical model representation in phylogenetics. Systematic biology. 2014;63(5):753–771.

45. du Plessis L, Stadler T. Getting to the root of epidemic spread with phylodynamic analysis of genomic data. Trends in Microbiology. 2015;23(7):383–386.

46. Tay JH, Baele G, Duchene S. Detecting episodic evolution through Bayesian inference of molecular clock models. Molecular Biology and Evolution. 2023;40(10):msad212.

47. Duchêne S, Holt KE, Weill FX, Le Hello S, Hawkey J, Edwards DJ, et al. Genome-scale rates of evolutionary change in bacteria. Microbial genomics. 2016;2(11):e000094.

48. Bouckaert R, Vaughan TG, Barido-Sottani J, Duchêne S, Fourment M, Gavryushkina A, et al. BEAST 2.5: An advanced software platform for Bayesian evolutionary analysis. PLoS computational biology. 2019;15(4):e1006650.

49. Ho SY, Shapiro B. Skyline-plot methods for estimating demographic history from nucleotide sequences. Molecular ecology resources. 2011;11(3):423–434.

50. Volz EM. Complex population dynamics and the coalescent under neutrality. Genetics. 2012;190(1):187–201.

51. Dearlove B, Wilson DJ. Coalescent inference for infectious disease: meta-analysis of hepatitis C. Philosophical Transactions of the Royal Society B: Biological Sciences. 2013;368(1614):20120314.

52. Wesner JS, Pomeranz JP. Choosing priors in Bayesian ecological models by simulating from the prior predictive distribution. Ecosphere. 2021;12(9):e03739.

53. Nascimento FF, Reis Md, Yang Z. A biologist’s guide to Bayesian phylogenetic analysis. Nature ecology & evolution. 2017;1(10):1446–1454.

54. Bouckaert RR. Tree priors and dating; 2021. Available from: beast2.blogs.auckland.ac.nz/tree-priors-and-dating/.

55. Rosenberg NA, Nordborg M. Genealogical trees, coalescent theory and the analysis of genetic polymorphisms. Nature Reviews Genetics. 2002;3(5):380–390.

56. Bouckaert RR. DensiTree: making sense of sets of phylogenetic trees. Bioinformatics. 2010;26(10):1372–1373.

57. Sanjuán R. From molecular genetics to phylodynamics: evolutionary relevance of mutation rates across viruses. PLoS pathogens. 2012;8(5):e1002685.

58. Baele G, Lemey P. Bayesian model selection in phylogenetics and genealogy-based population genetics. In: Chen MLKPOL, editors. Bayesian phylogenetics, methods, algorithms, and applications. Boca Raton (Florida): CPC Press; 2014. p. 59–93.

59. Boskova V, Stadler T, Magnus C. The influence of phylodynamic model specifications on parameter estimates of the Zika virus epidemic. Virus evolution. 2018;4(1):vex044.

60. Bergsten J, Nilsson AN, Ronquist F. Bayesian tests of topology hypotheses with an example from diving beetles. Systematic biology. 2013;62(5):660–673.

61. Warnock RC, Parham JF, Joyce WG, Lyson TR, Donoghue PC. Calibration uncertainty in molecular dating analyses: there is no substitute for the prior evaluation of time priors. Proceedings of the Royal Society B: Biological Sciences. 2015;282(1798):20141013.

62. Foster CS, Sauquet H, Van der Merwe M, McPherson H, Rossetto M, Ho SY. Evaluating the impact of genomic data and priors on Bayesian estimates of the angiosperm evolutionary timescale. Systematic Biology. 2017;66(3):338–351.

63. Lopes HF, Tobias JL. Confronting prior convictions: On issues of prior sensitivity and likelihood robustness in Bayesian analysis. Annu Rev Econ. 2011;3(1):107–131.

64. Lambert B. A student’s guide to Bayesian statistics. London: SAGE Publications Ltd; 2018.

65. Duchene S, Duchene DA. Estimating evolutionary rates and timescales from time-stamped data. The Molecular Evolutionary Clock: Theory and Practice. 2020; p. 157–174.

66. Duchêne S, Lanfear R, Ho SY. The impact of calibration and clock-model choice on molecular estimates of divergence times. Molecular phylogenetics and evolution. 2014;78:277–289.

67. Duchene S, Featherstone L, Haritopoulou-Sinanidou M, Rambaut A, Lemey P, Baele G. Temporal signal and the phylodynamic threshold of SARS-CoV-2. Virus Evolution. 2020;6(2):veaa061.

68. Baele G, Lemey P, Suchard MA. Genealogical working distributions for Bayesian model testing with phylogenetic uncertainty. Systematic Biology. 2016;65(2):250–264.

69. Fan Y, Wu R, Chen MH, Kuo L, Lewis PO. Choosing among partition models in Bayesian phylogenetics. Molecular Biology and Evolution. 2011;28(1):523–532.

70. Porter AF, Sherry N, Andersson P, Johnson SA, Duchene S, Howden BP. New rules for genomics-informed COVID-19 responses–lessons learned from the first waves of the omicron variant in Australia. PLoS Genetics. 2022;18(10):e1010415.

71. Zhang C, Stadler T, Klopfstein S, Heath TA, Ronquist F. Total-evidence dating under the fossilized birth–death process. Systematic biology. 2016;65(2):228–249.

72. Stadler T, Vaughan TG, Gavryushkin A, Guindon S, Kühnert D, Leventhal GE, et al. How well can the exponential-growth coalescent approximate constant-rate birth–death population dynamics? Proceedings of the Royal Society B: Biological Sciences. 2015;282(1806):20150420.

73. Volz EM, Frost SD. Sampling through time and phylodynamic inference with coalescent and birth–death models. Journal of The Royal Society Interface. 2014;11(101):20140945.

74. Jukes TH, Cantor CR, et al. Evolution of protein molecules. Mammalian protein metabolism. 1969;3(24):21–132.

75. Ho SY, Duchêne S, Duchêne D. Simulating and detecting autocorrelation of molecular evolutionary rates among lineages. Molecular ecology resources. 2015;15(4):688–696.

76. Schliep KP. phangorn: phylogenetic analysis in R. Bioinformatics. 2011;27(4):592–593.

